# System identification and surrogate data analyses imply approximate Gaussianity and non-stationarity of resting-brain dynamics

**DOI:** 10.64898/2026.03.25.714361

**Authors:** Teppei Matsui, Ruixiang Li, Koki Masaoka, Koji Jimura

## Abstract

Compared with model-based and phenomenological descriptions of the spatiotemporal dynamics of resting-brain activity, statistical characterizations of resting-state fMRI (rs-fMRI) data remain relatively underexplored. Some sophisticated analysis techniques, such as Mapper-based topological data analysis (TDA) and innovation-driven coactivation pattern analysis (iCAP), can distinguish real data from phase-randomized (PR) surrogates, suggesting that rs-fMRI data are not as simple as stationary Gaussian processes. However, the exact statistical properties that distinguish real rs-fMRI data from PR surrogates have not yet been determined. In this study, we conducted system identification analysis and surrogate data analysis to specify key statistical properties that allow TDA and iCAP to discriminate real rs-fMRI data from PR surrogates. We first analyzed rs-fMRI data concatenated across scans using autoregressive (AR) modeling and found that the scan-concatenated rs-fMRI data were weakly non-Gaussian. However, non-Gaussianity alone was insufficient to reproduce realistic TDA and iCAP results because of non-stationarity across scans. AR modeling of single-scan data revealed that rs-fMRI data were statistically indistinguishable from a Gaussian distribution within a single scan, although TDA and iCAP results still differed between the real data and PR surrogates. A new surrogate dataset designed to preserve non-stationarity successfully reproduced realistic TDA and iCAP results, suggesting that TDA and iCAP likely capture the non-stationarity of rs-fMRI data to distinguish it from PR surrogates. Together, these results indicate approximate Gaussianity and non-stationarity in rs-fMRI data, providing a data-driven and statistical characterization of resting-state brain activity that can serve as a quantitative reference for whole brain simulations and generative models.

## Introduction

Spontaneous brain activity exhibits rich spatiotemporal dynamics that have been extensively investigated, particularly in the field of resting-state functional magnetic resonance imaging (rs-fMRI) (*1–10*). Conceptually, rs-fMRI data are commonly modeled as non-stationary dynamics, such as transitions among locally stable brain states (*9, 11–14*). However, the extent to which this assumption is supported by empirical rs-fMRI data has remained an important yet controversial research topic (*1, 2, 12, 15–22*). Although many techniques have been proposed to extract non-stationary dynamical features from rs-fMRI data, *e.g.,* putative brain states extracted by sliding-window correlation (*1*) and coactivation pattern analysis (*23*), other studies have demonstrated that these dynamical features and putative brain states can be reproduced using surrogate data generated by autoregressive (AR) modeling or phase randomization (PR) (*16, 17, 24–27*). Moreover, AR surrogates also reproduce standing and traveling waves in the real data (*5*). Because AR and PR surrogates are stationary, linear, and Gaussian by construction, these findings suggest that, at least as a first approximation, rs-fMRI dynamics may be simple enough to be modeled as linear, stationary, and Gaussian processes (*15, 28*).

However, considering previous electrophysiological studies reporting non-stationary and non-Gaussian spontaneous brain activity (*29–32*), the statement that rs-fMRI data are approximately linear, stationary, and Gaussian appears counterintuitive. Notably, recent studies employing advanced analytical techniques have suggested that rs-fMRI data are indeed more complex. Studies using Mapper-based topological data analysis (TDA) and innovation-driven co-activation pattern analysis (iCAP) demonstrated that these methods can distinguish real rs-fMRI data from PR surrogates (*13, 33*). According to these studies, rs-fMRI data must therefore be non-Gaussian, non-stationary, non-linear, or some combination of these properties (*21*).

While these studies significantly advanced our understanding of rs-fMRI dynamics, the specific statistical properties that enable TDA and iCAP to differentiate real data from surrogates remain unidentified. To address this question, we performed a series of system identification and surrogate data analyses to identify the key statistical properties characterizing rs-fMRI data (Fig. 1). We first conducted system identification analyses based on AR modeling to capture deterministic temporal structures in rs-fMRI data (Fig. 1, left). By analyzing the residuals of the fitted AR models, we further assessed the Gaussianity and stationarity of the data. We also conducted surrogate data analyses (Fig. 1, right), in which various surrogate datasets retaining specified sets of real statistics were generated. TDA and iCAP were applied to both the real and surrogate data to identify the key statistical properties that distinguish real data from AR and PR surrogates.

**Figure 1.**
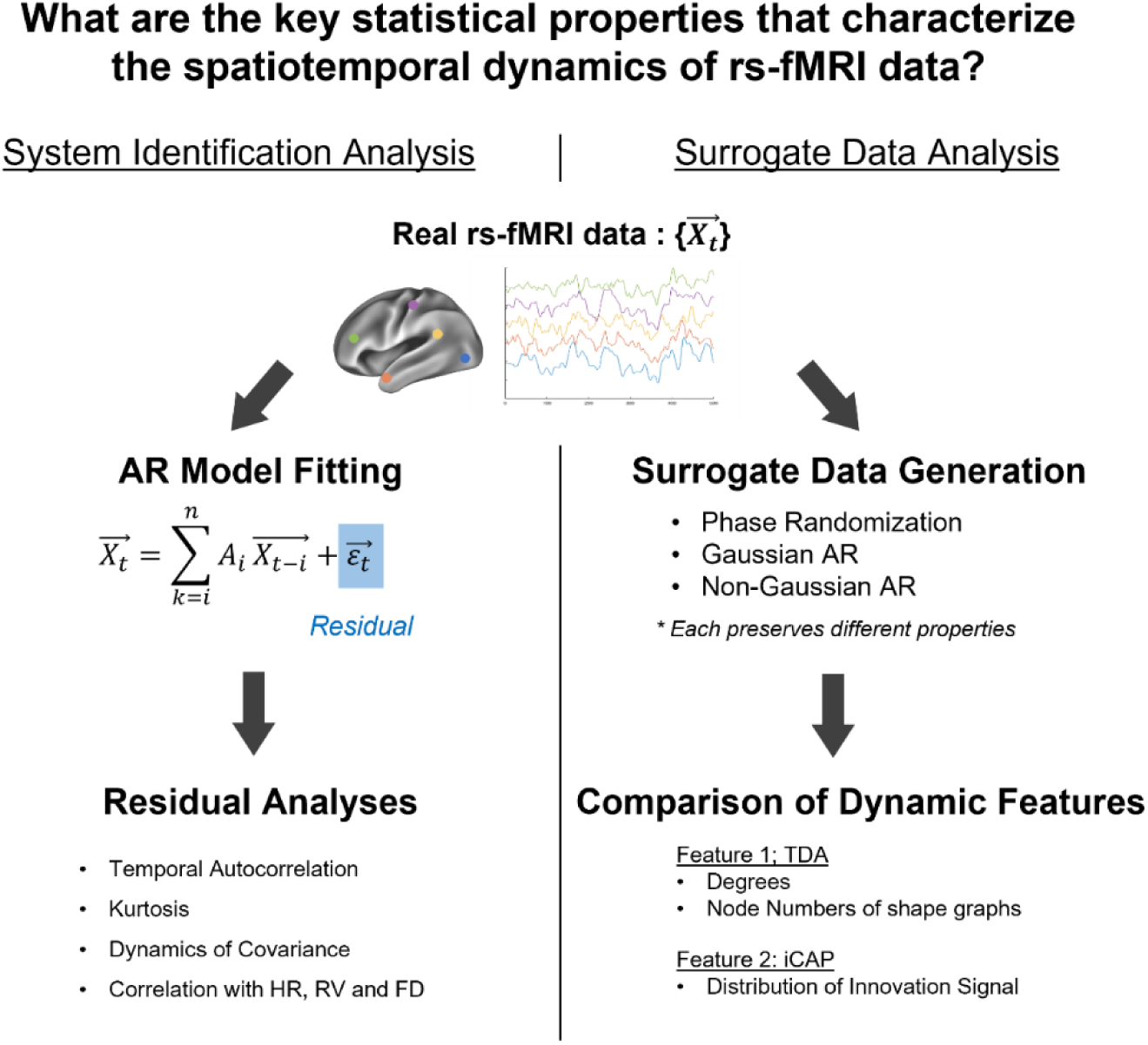
Overview of the study. System Indentification and surrogate data analyses were performed using rs-fMRI data from Human Connectome Project (HCP). **Left:** Procedures for system identification analysis. Rs-fMRI data were fitted by an AR model of order *n*. Residual analyses were then performed to examine temporal autocorrelation (Fig. 2E, Fig. 4E), kurtosis (Fig. 2F,G, Fig. 4F,G), independent components (Fig. 4) and correlations with physiological and behavioral variables (Fig. 6). **Right:** Procedures for surrogate data analysis. For each real rs-fMRI dataset, surrogate data of a certain type were generated. Each surrogate dataset preserved selected statistical properties (e.g., covariance) of the real data. TDA and iCAP were then applied to both the real and surrogate datasets, and the extracted dynamical features were compared (Fig. 2A-D, Fig. 4A-D; Fig. 5). TDA and iCAP were chosen because of their demonstrated ability to distinguish the real data from and PR surrogates.

## Results

### TDA and iCAP can distinguish real rs-fMRI data from PR surrogates

Our goal was to understand statistical properties of rs-fMRI data. Specifically, we aimed to identify the statistical properties that allow TDA and iCAP to discriminate real rs-fMRI data from PR surrogates. We used rs-fMRI data provided by Human Connectome Project (HCP), preprocessed with a standard preprocessing pipeline for rs-fMRI, including motion scrubbing, band-pass temporal filtering, and parcellation using the Schaefer100 atlas (*34*) (see Methods for details).

In the first set of analyses (Figs. 2 and 3), we harmonized and concatenated the four rs-fMRI scans for each participant, as commonly done in previous studies. Consistent with previous TDA and iCAP studies, both TDA and iCAP results differed significantly between the real data and PR surrogates (Fig. 2A-D). For TDA, as reported previously (*33*), the fraction of high-degree nodes was significantly higher in the real data than in the PR surrogates (p < 0.001, post-hoc Tukey’s test) (green trace, Fig. 2A). Additionally, the number of nodes in the shape graphs was significantly lower for the real data than for the PR surrogates (p < 0.05, post-hoc Dunn’s test corrected by Bonferroni’s method) (Fig. 2B). For iCAP, the distributions of the innovation signal obtained from the real data were significantly more heavy-tailed than those obtained from PR surrogates (p < 0.001, post-hoc Dunn’s test corrected by Bonferroni’s method) (green trace, Fig. 2C-D), consistent with previous findings (*13*). These results confirm previous TDA and iCAP studies and suggest that rs-fMRI data are non-Gaussian, non-linear, non-stationary or some combination of these properties.

**Figure 2.**
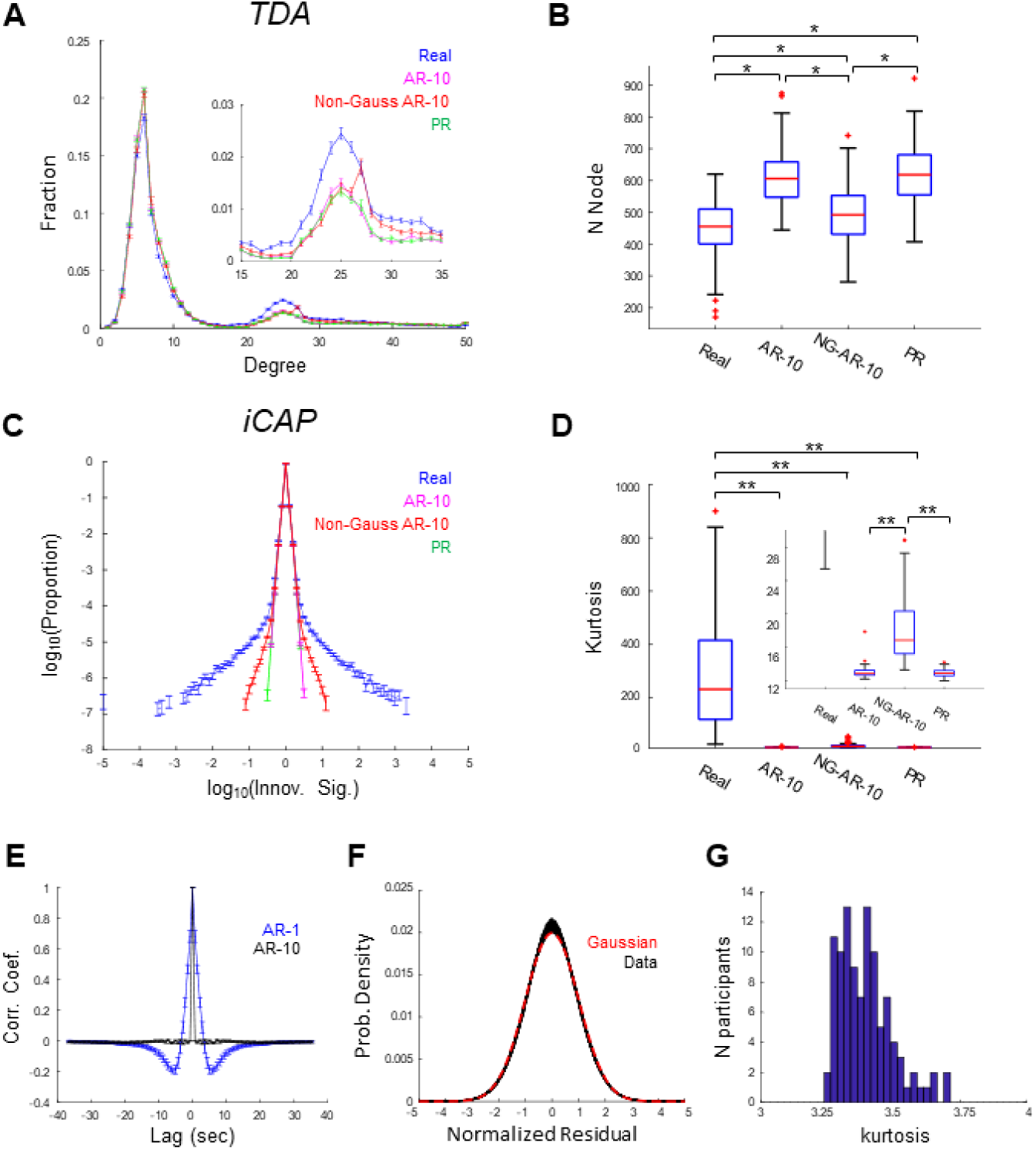
TDA and iCAP applied to scan-concatenated data. Non-Gaussianity alone does not explain discrepancy of TDA and iCAP results between real and surrogate data. **(A)-(B)** TDA-based features for real and surrogate data. (A) Degree distributions of the shape-graphs. Inset: enlarged view showing a prominent difference between real and surrogate data. Error bar: SE. (B) Boxplots of numbers of nodes of the shape-graphs. **(C)-(D)** iCAP-based features for real and surrogate data. (C) Distribution of the innovation signal. Error bar: SE. (D) Boxplots of kurtosis of the distribution of innovation signal. Inset: enlarged view showing difference between surrogates. **(E)** Autocorrelation functions of the residuals of AR-1 (blue) and AR-10 (black). **(F)** Distribution of the normalized residuals. Black traces: individual participants (n = 103). Red trace: Gaussian distribution. **(G)** Distribution of the kurtosis of the normalized residual across participants. *, p < 0.05, **, p < 0.001 (Dunn’s test corrected by Bonferroni’s method).

**Figure 3.**
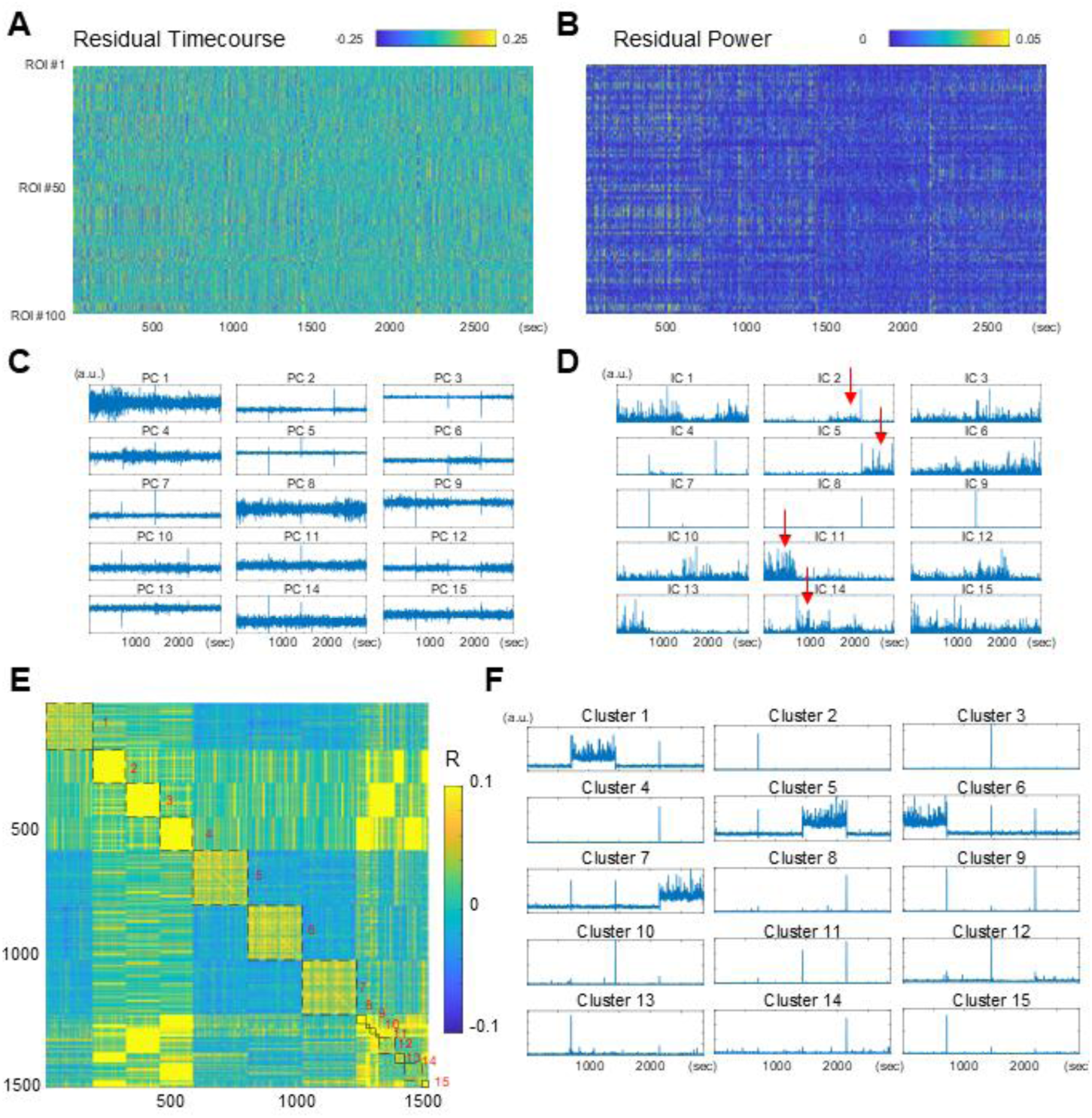
AR-10 residuals of scan-concatenated data exhibited non-stationarity. **(A)** Example residual timecourse. **(B)** Example timecourse of residual power. Same data as in (A). **(C)** PCs of the residual timecourse. Same data as in (A). **(D)** Power timecourses of ICs of the residual timecourse. Same data as in (A). Red arrows, ICs capturing the structure of fMRI scans. **(E)** Correlation between the power timecourses of all ICs in all participants (1545 ICs). Sorted by k-means clustering (15 clusters). **(F)** Average power timecourse for each cluster of ICs.

### Scan-concatenated rs-fMRI data were weakly non-Gaussian

To examine Gaussianity, we next fitted AR models to rs-fMRI data to achieve temporal decorrelation. We first fitted the first order AR model (AR-1), which has been shown to account for the dynamics of the rs-fMRI data reasonably well (*17, 35*), and analyzed its residuals. However, we found that the residuals of the AR-1 showed strong temporal autocorrelation (Fig. 1A), suggesting that AR-1 failed to account for all linear and deterministic temporal structures in the data (*36*). To fully account for the linear, deterministic component and temporally decorrelate the rs-fMRI data, we increased the order of the AR model. We found that the autocorrelation functions of the residuals became close to a delta function when the model order was increased to ten (AR-10) (Fig. 2A), suggesting that AR-10 achieved temporal decorrelation of the rs-fMRI data.

To further remove spatial correlation, principal component analysis (PCA) was applied to the residuals of the AR-10 model (AR-10 residuals). The signal of each component was then standardized and concatenated to examine its Gaussianity (normalized residuals). The distributions of the normalized residuals were well fit by Pearson distribution functions with kurtosis of approximately 3.4 (Fig. 2E-F), indicating slightly heavier tails than the Gaussian distribution (kurtosis = 3; Fig. 2F, red). Although the deviation from the Gaussian was small, all individual distributions of normalized residuals were significantly difference from the Gaussian distribution (n = 103 participants; p < 0.01, KS test, uncorrected) (Fig. 2G). The distributions of the normalized residuals calculated from PR surrogates were approximately Gaussian (kurtosis = 3.0; p> 0.01 for all 103 participants, KS test, uncorrected) (Fig. S1A). These results suggest that scan-concatenated rs-fMRI data are weakly but significantly non-Gaussian.

### Non-Gaussianity alone did not fully explain the discrepancies between scan-concatenated rs-fMRI data and PR surrogates

We next asked whether non-Gaussianity alone was sufficient to reproduce realistic TDA and iCAP results. To this end, non-Gaussian surrogate data were generated by temporally shuffling the AR-10 residuals (non-Gaussian AR surrogates). If TDA and/or iCAP produce significantly different results for real rs-fMRI data and non-Gaussian AR surrogates, this would imply that rs-fMRI data are non-stationary and/or nonlinear.

We found that TDA and iCAP results obtained with the non-Gaussian AR surrogates were still different from those obtained with the real data. For TDA, we found that high-degree nodes were more abundant in shape graphs derived from the real data than from Gaussian surrogates (Gaussian AR and PR) and non-Gaussian AR surrogates (p < 0.001, post-hoc Tukey’s test; red trace, Fig. 2A inset). Similarly, the total number of nodes in the shape graphs was significantly lower for real data than for both Gaussian and non-Gaussian surrogates (p < 0.001, post-hoc Dunn’s test corrected by Bonferroni’s method; Fig. 2B). The results for the non-Gaussian AR surrogates were positioned between those of the Gaussian surrogates and the real data, suggesting that accounting for non-Gaussianity partially reduced the discrepancy between the surrogates and the real data.

For iCAP, the distribution of the innovation signal was significantly more heavy-tailed in the real data than in both Gaussian and non-Gaussian surrogates (p < 0.001, post-hoc Dunn’s test corrected by Bonferroni’s method; Fig. 2C-D). Again, the distributions of the non-Gaussian surrogates had significantly heavier tail than those of the Gaussian surrogates (red trace, Fig. 2C inset), but they still failed to match those of the real data. Taken together, these results suggest that weak non-Gaussianity alone cannot account for the observed differences, and that rs-fMRI data must also exhibit non-stationarity and/or nonlinearity.

### rs-fMRI data are non-stationary across scans

Because non-Gaussianity alone was insufficient, we next examined possible non-stationarity of rs-fMRI data by analyzing the AR-10 residuals (Fig. 3A). Visual inspection of the power of the residuals (Fig. 3B), as well as of their principal components (PC) (Fig. 3C), revealed block-like structures reminiscent of concatenated fMRI scans. A similar structure was also found in the power of some of the independent components (ICs) (Fig. 3D, arrows).

If these block-like structures indeed reflect concatenated fMRI scans, then such structures should be present across all participants. To verify this point, we computed ICs of the residuals for all participants and clustered them based on the similarity of their power time courses (Fig. 3E, left), and then averaged the power time courses within each cluster (Fig. 3E, right). Many of the resulting power time courses clearly represented individual fMRI scans or the transitions between them. The presence of these ICs indicates that, despite harmonization during preprocessing, rs-fMRI data exhibit substantial non-stationarity across different scans. The difference between real rs-fMRI data and PR surrogates in TDA and iCAP likely reflect this across-scan non-stationarity.

### rs-fMRI data were approximately Gaussian within single scans

The non-Gaussianity of scan-concatenated rs-fMRI data may arise from across-scan non-stationarity. Hence, we next analyzed single-scan rs-fMRI data to clarify these points. We first confirmed that TDA and iCAP results for single-scan data were significantly different between the real data and PR surrogates (Fig. 4A-D). In TDA, high-degree nodes were more abundant in shape graphs derived from the real data than from the surrogate data (p < 0.001, post-hoc Tukey’s test; Fig. 4A, inset). Moreover, the total number of nodes in the shape graphs was significantly lower for the real data than for the surrogate data (p < 0.001, post-hoc Dunn’s test corrected by Bonferroni’s method; Fig. 4B). In iCAP, the innovation signal distributions were significantly more heavy-tailed in the real data than in the surrogate data (p < 0.001, post-hoc Dunn’s test corrected by Bonferroni’s method; Fig. 4C-D).

**Figure 4.**
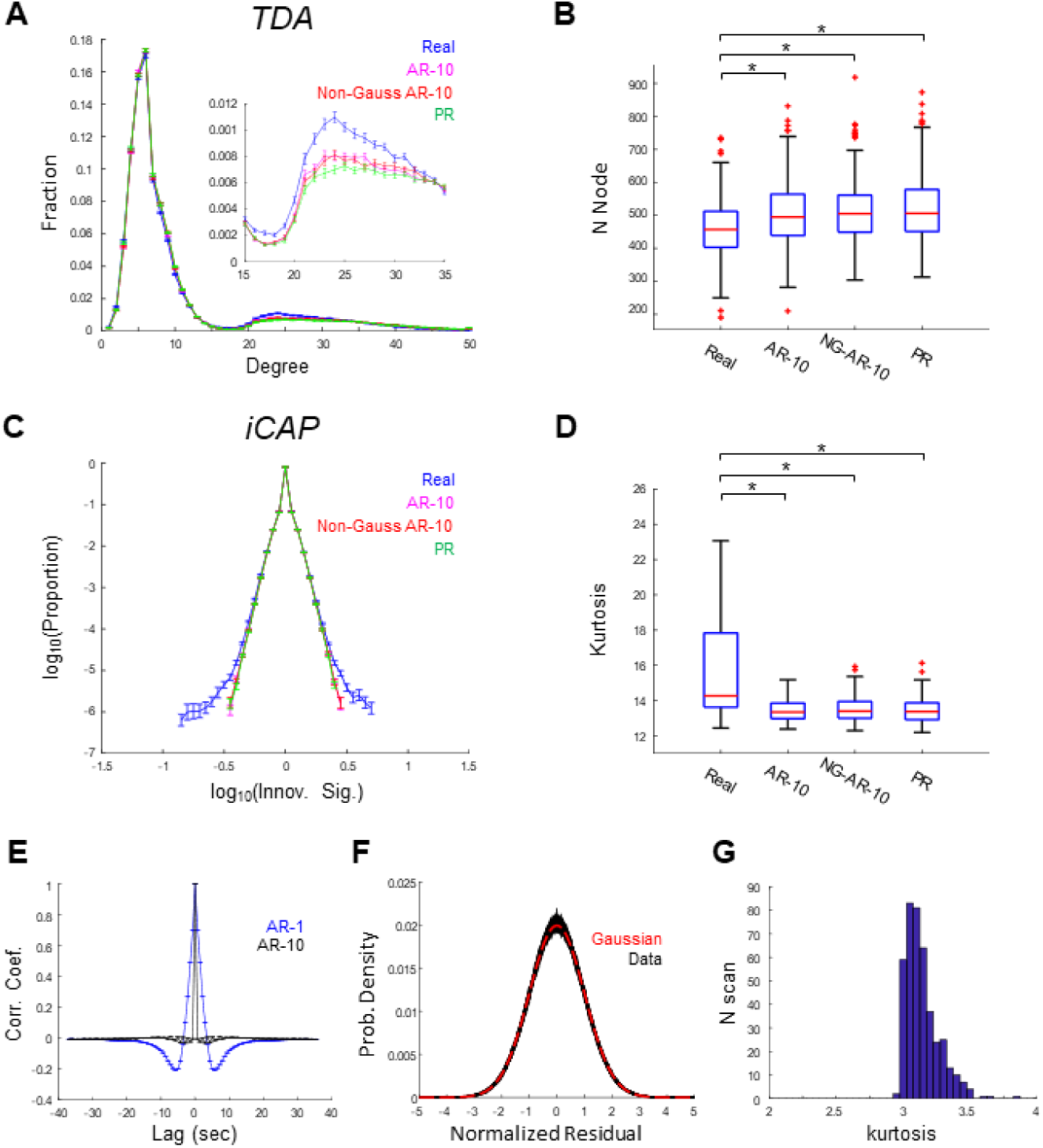
TDA and iCAP applied to single-scan data. **(A)-(B)** TDA-based features for real and surrogate data. (A) Degree distributions of shape-graphs. Inset: enlarged view showing the prominent difference between real and surrogate data. Error bars indicate the standard error (SE). (B) Boxplots showing the numbers of nodes in shape-graphs. **(C)-(D)** iCAP-based features for real and surrogate data. (C) Distribution of the innovation signal. Error bars indicate SE. (D) Boxplots of kurtosis values for the distribution of the innovation signal. **(E)** Autocorrelation functions of the residuals from AR-1 (blue) and AR-10 (black) models. **(F)** Distribution of normalized residuals, normalized to zero mean and unit variance. Black traces indicate individual scans (n = 412); the red trace shows a Gaussian distribution. **(G)** Distribution of the kurtosis values of the normalized residuals across scans. *, p < 0.001 (Dunn’s test, Bonferroni corrected).

To examine possible non-Gaussianity, we fitted AR models to single-scan rs-fMRI data and analyzed the residuals in detail. Similar to the scan-concatenated data, the residuals from AR-1 models exhibited substantial temporal autocorrelation (Fig. 4E, blue). Thus, we used AR-10 models to fully separate the linear and deterministic components from the stochastic component in the data. To fit the AR-10 model with single-scan data, we used a coarse cortical parcellation with 34 parcels (*34*). The temporal autocorrelation function of the AR-10 residuals resembled a delta function (Fig. 4E, black), suggesting that the AR-10 model accounted for most of the linear and deterministic temporal structures.

We then examined the Gaussianity of the AR-10 residuals by fitting Pearson distribution functions to the normalized residuals (Fig. 4F). The Gaussian distribution fell within the range of the best-fitting Pearson distribution functions obtained from the data, suggesting that the normalized residuals were approximately Gaussian (kurtosis = 3.1; Fig. 4F-G). Indeed, the distributions of the normalized residuals in most of the scans (382/412) did not significantly differ from the Gaussian distribution [p < 0.01, ks-test (uncorrected), in 30/412 scans]. Consistently, TDA and iCAP results did not significantly differ between non-Gaussian AR-10 surrogates and Gaussian AR-10 surrogates (Fig. 4A-D). The distributions of the normalized residuals calculated from PR surrogates were approximately Gaussian (kurtosis = 3.0; p> 0.01 for all 412 participants, KS test, uncorrected) (Fig. S1B). These results suggest that, within each scan, rs-fMRI data are approximately Gaussian. Moreover, Gaussianity of the data suggests that the differences between the real data and the surrogates must be explained by non-stationarity and/or nonlinearity.

### Non-stationarity accounted for realistic TDA and iCAP results

To examine whether non-stationarity was sufficient to reproduce realistic TDA and iCAP results, we generated novel non-stationary surrogate data by applying a block-wise shuffling procedure to the AR-10 residuals. The block-shuffling approach divides a scan into smaller temporal segments and permutes their order while preserving temporal dependencies within each segment (Fig. 5A), unlike frame-wise shuffling, which completely destroys temporal structure. We reasoned that the block shuffling would randomize the data without disrupting potential temporal dependency in the data, provided that the block size was sufficiently larger than the extent of the temporal dependency.

**Figure 5.**
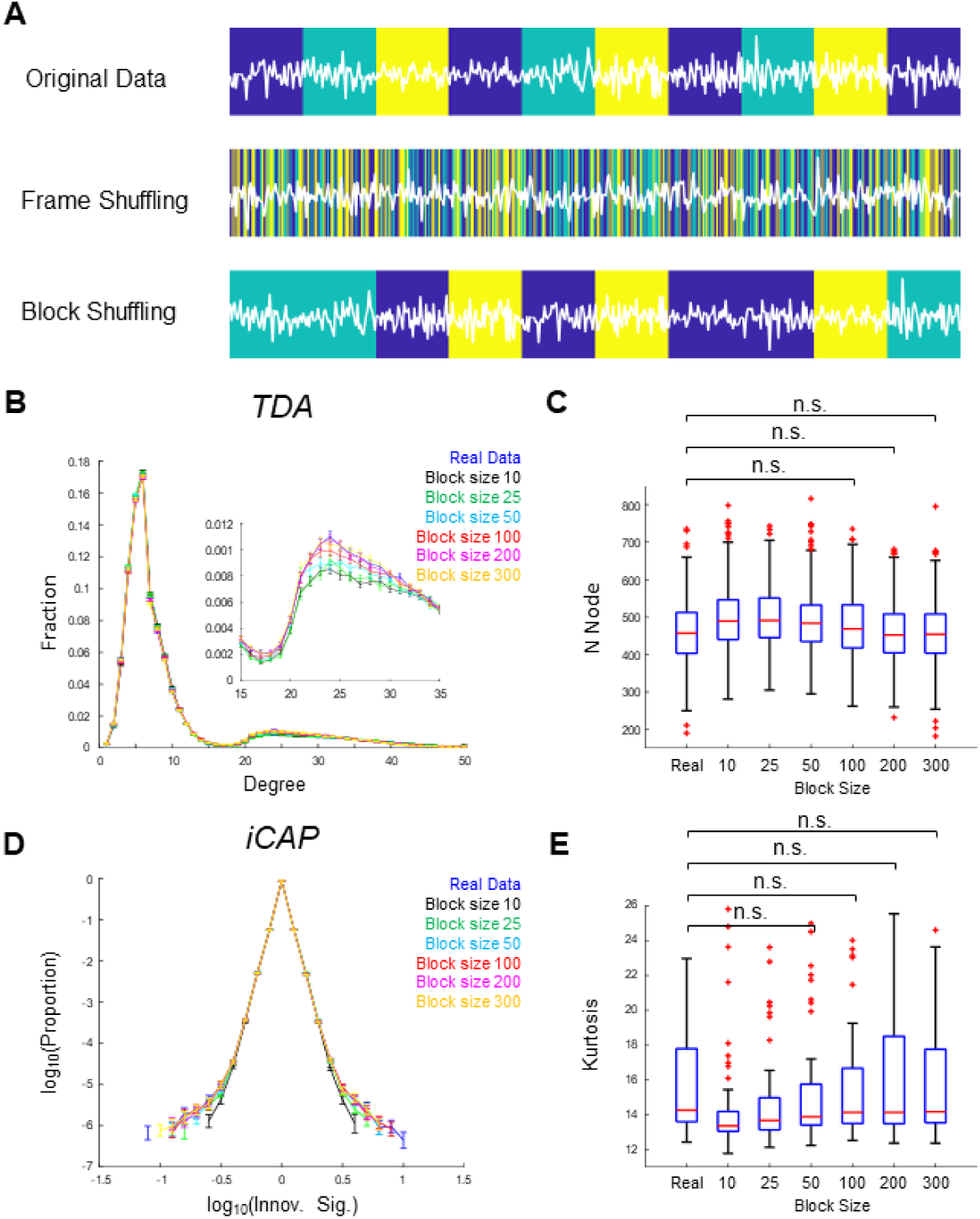
Surrogate data incorporating non-stationarity successfully recapitulated TDA- and iCAP-based features of real data. **(A)** Schematic illustration of block shuffling. Original data (top) alternate between states represented by colored blocks. Frame shuffling (middle) destroys state information, whereas block shuffling preserves it when the block size is sufficiently large. **(B)-(C)** TDA-based features for real and surrogate data. (B) Degree distributions of shape-graphs. Error bars indicate SE. (C) Boxplots showing the numbers of nodes in shape-graphs. No significant differences were observed between real data and surrogated with block sizes ≥ 100 TRs. **(D)-(E)** iCAP-based features for real and surrogate data. (D) Distribution of the innovation signal. Error bars indicate SE. (E) Boxplots of kurtosis values for the distribution of the innovation signal. No significant differences were observed between real and surrogate data with block sizes ≥ 50 TRs. p > 0.05 (Dunn’s test, Bonferroni corrected). See Fig S2 for significant differences.

Consistent with our expectation, block-shuffling produced surrogate data that successfully replicated the TDA and iCAP results obtained from the real data. When the block size exceeded 100 TRs (72 s), no statistically significant differences were observed between real and surrogate data for either TDA or iCAP (p > 0.05, post hoc Dunn’s test corrected by Bonferroni’s method) (Fig. 5B-E) (See Fig. S2 for significant differences). These findings suggest that real rs-fMRI data were approximately Gaussian but non-stationary, with a characteristic timescale of approximately 70 s, and that this non-stationarity was likely captured by TDA and iCAP.

### Non-stationarity of resting-brain activity correlated with arousal-related variables

So far, we have analyzed the AR-10 residuals to characterize the Gaussianity and non-stationarity of rs-fMRI data. However, because an AR-10 model fits the data with a large number of parameters, one concern was that the AR-10 residuals might primarily reflect overfitting rather than biological activity. Alternatively, the AR-10 residuals could reflect measurement errors (*37*) or sampling errors (*16*). To address these concerns, we next examined potential biological relevance of the AR-10 residuals.

Based on recent studies showing that spontaneous brain activity in the resting-state is modulated by physiological states of the body (*38, 39*) and behavioral factors (*40, 41*), we examined the relationship between the first principal component of the power of the AR-10 residuals (hereafter referred to as *residual power*) and three non-neuronal physiological/behavioral variables: heart rate (HR), respiration variation (RV) (*42*), and framewise displacement (FD), which serves as an indicator of head-motion (*43*). Although the preprocessing of rs-fMRI data in the present study included FD-based frame censoring, we still assessed the relationship between residual power and FD, motivated by recent reports suggesting that a substantial fraction of spontaneous neuronal activity reflects body movements (*40, 41*).

As shown in representative examples, both HR and RV exhibited moderate but statistically significant correlations with residual power (Fig. 6A, top and middle rows). FD showed a markedly higher correlation with residual power (Fig. 6A, bottom row), which was unexpected given the extensive preprocessing to remove motion-related artifacts (*44*). At the population level, the correlations of residual power with both HR and RV were significantly higher than those in the shuffled control, even after partializing out the effect of FD (p < 0.001 and p < 0.05 by two-sample t test, for HR and RV, respectively; Fig. 6B-C). Importantly, significant differences from the control persisted even when the analyses were restricted to volumes with small FD values (Fig. S2), suggesting that the observed correlations between residual power and physiological variables were unlikely to be driven by motion-related artifacts. Taken together, these results indicate that the AR-10 residuals are unlikely to be artefacts produced by overfitting of the AR-10 model; rather, they reflect biological brain states coupled to arousal levels.

**Figure 6.**
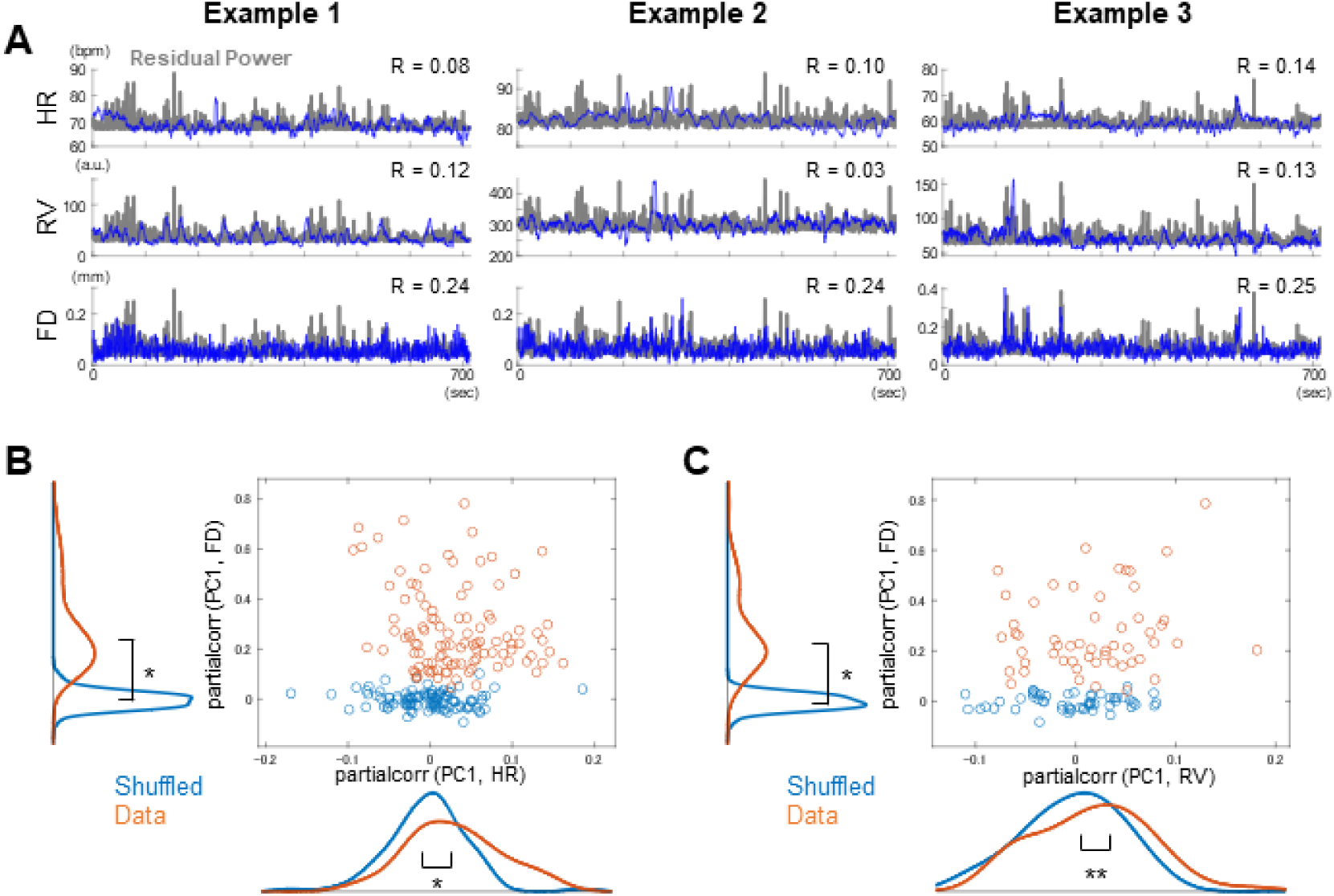
AR-10 residuals correlated with physiological and behavioral variables. **(A)** Example plots showing the relationship between the residual amplitude and non-neuronal variables. Blue traces indicate time courses of HR, RV, FD. Gray traces indicate time courses of the PC1 of the power of the residual. R indicates correlation between each variable and the residual power. **(B)** Scatter plot showing correlations between PC1 and HR partialized for FD, and between PC1 and FD partialized for HR, for 104 scans with reliable HR measurements. Orange, real data; Blue, shuffled data. Marginal smoothed histograms are shown along the top and left. **(C)** Scatter plot showing correlation between PC1 and RV partialized for FD, and between PC1 and FR partialized for HR, for 51 scans with reliable RV measurements. Orange, real data; Blue, shuffled data. Marginal smoothed histograms are shown along the top and left. *, p < 0.001, **, p < 0.05 (two-sample t-test).

## Discussion

In the present study, we performed AR-based system identification and a series of surrogate data analyses to determine the statistical properties that characterize the spatiotemporal dynamics of rs-fMRI data. The residuals from the AR-10 model fitted to single-scan rs-fMRI data were approximately Gaussian but exhibited non-stationarity. Surrogate data incorporating non-stationarity successfully reproduced realistic TDA and iCAP results. Together, these results demonstrate that 1) rs-fMRI data are approximately Gaussian within each scan; 2) the spatiotemporal dynamics of rs-fMRI data are non-stationary within each scan and even more so across scans. Furthermore, the correlation between the amplitude of the residuals and physiological and behavioral variables suggests that the non-stationarity of single-scan rs-fMRI data is likely coupled to fluctuations in arousal levels.

Recently, it has been argued that many dynamic features of rs-fMRI data can be reproduced using linear, stationary, and Gaussian surrogates (e.g., PR surrogates) (*5, 16, 17, 24–26, 35*). Notable exceptions are the dynamic features obtained by TDA and iCAP, which cannot be reproduced by the PR surrogates (*13, 33*). The present study extends these findings by showing that TDA and iCAP can detect non-stationarity inherent in rs-fMRI data. Note that TDA and iCAP were used to detect differences between the real and surrogate data, and neuroscientific interpretations of the TDA and iCAP results (e.g., graph structures or brain-states) were outside of the scope of this study. Interestingly, our results also indicate that nonlinear models are not required to reproduce these features, consistent with recent studies reporting approximately linear dynamics of rs-fMRI signals (*35, 45*). It is important to note, however, that our findings merely suggest that TDA and iCAP do not necessitate nonlinear dynamics at the level modeled here. Future studies comparing linear and nonlinear models, such as the ordinary differential equation-based models (*46*), will be necessary to clarify the degree of nonlinearity present in rs-fMRI dynamics.

The present results suggest that rs-fMRI data are approximately Gaussian within each fMRI scan. Even in scan-concatenated data, deviation from Gaussianity were subtle. These observations are consistent with earlier fMRI studies showing that the distribution of rs-fMRI data is largely Gaussian (*47, 48*). The approximate Gaussianity is also consistent with the observation that Gaussian surrogate data can reproduce many features of rs-fMRI dynamics (*16, 17, 24*). In contrast to these observations, non-Gaussian, heavy-tailed distributions of spontaneous brain activity have previously been reported in electrophysiological studies (*29, 30*). Elucidating the precise mechanisms underlying the emergence of Gaussianity in rs-fMRI signals will be an interesting question for future studies.

We also found that rs-fMRI data are non-stationary both within individual scans and across concatenated scans. These findings are consistent with previous studies reporting non-stationarity in functional connectivity (*8, 49*). Notably, strong non-stationarity across multiple fMRI scans persisted despite the use of harmonization procedures. This observation has practical implications, as it is common to combine multiple rs-fMRI scans from the same participant to increase statistical power or to fit large-scale models (*33, 50, 51*). It should be noted that some components of across-scan non-stationarity may have non-neuronal origins. For instance, a previous study reported that respiration-related artifacts in rs-fMRI signals depend on the phase-encoding direction of MRI acquisition (*52*).

Weaker but significant non-stationarity within individual rs-fMRI scans was also observed. This finding is consistent with a previous rs-fMRI study reporting non-stationary functional connectivity dynamics in the default mode network (*8*), as well as a previous calcium imaging study in mice conducted by our group (*27*). The present finding is largely in agreement with these studies. Moreover, we found that the amplitude of resting-brain activity, was significantly correlated with the non-neuronal physiological measures as well as with the behavioral measure. These correlations align with recent studies suggesting that spontaneous brain activity is tightly coupled to the animal’s bodily states and behavior (*38–41*). The strong correlation between the residual power and head-motion as quantified by FD (Fig. 6) was surprising, given careful removal of motion-related artefacts in the preprocessing. A control analysis restricted to volumes with small head-motion (Fig. S3) suggest that this correlation is unlikely to be simply driven by motion-related artefacts. Nevertheless, given the strength of correlation, there is a possibility that remaining motion-related artefacts are affecting the correlation between the residual power and FD. Such non-biological artifacts likely contributed to at least part of the within-scan and across-scan non-stationarity observed in the present study. Future animal studies with calcium imaging would be useful to further address these points (*53, 54*).

Whereas the present study took a bottom-up approach to characterize rs-fMRI data, the results obtained here may also be informative for top-down approaches, in particular generative models of resting-brain activity (*7, 12, 53, 55, 56*). First, quantitative and statistical characteristics of rs-fMRI data revealed in the present study could serve as constraints that a generative model must satisfy. Second, the present results suggest that rs-fMRI data can be can be described as a high-dimensional linear dynamical system driven by a non-stationary, arousal-driven process. This architecture is consistent with the proposal that an arousal-related nonlinear dynamical system, which can be lifted to linear high dimensional system, constitutes the principal generative mechanism underlying resting-state infra-slow brain activity (*53*). Future studies integrating bottom-up and top-down approaches will be essential for developing realistic generative models of resting-brain activity, which can in turn be used to better understand brain functioning in health and disease.

## Materials and Methods

### Dataset and preprocessing

We used the S1200 release of resting-state fMRI data distributed by the Human Connectome Project (http://humanconnectomeproject.org/)(1200 volumes × 64984 vertices × 1003 participants; repetition time (TR), 0.72 s). We used data processed using independent component analysis (ICA)-based denoising (ICA-FIX) (*57*). Cortical resting-state activity was obtained using Schaffer100 (100 cortical parcels)(*34*) and Yeo17 (34 cortical parcels)(*58*) for scan-concatenated data and single scan data, respectively. First, we first selected 103 participants with small head motions. The number of participants matched the sample size of a previous study using TDA (*33*). To quantify head motions, the framewise displacement (FD) was calculated from the rigid-body motion parameters. Notch filters (0.31 Hz < f < 0.43 Hz) were applied to the rigid-body motion parameters before computing FD to remove respiratory artifacts in head-motion estimates (*44*). The fMRI volumes were flagged when the FD exceeded 0.2. The volumes before and after the flagged volumes were also flagged. Flagged volumes were discarded, and participants with > 20% of discarded volume (∼30% of participants) were excluded from further analyses. The discarded volumes were replaced with linear interpolation of the uncensored volumes (*33*). After the motion-scrubbing, temporal bandpass filters retaining frequencies between 0.01 Hz and 0.1 Hz were applied, and the first and the last 100 volumes of each scan were discarded. Nuisance regressions were then performed using the six motion parameters, their derivatives, and the global signal (the global mean of the time course). To allow harmonization across scans, the time course of each parcel was normalized to have a zero mean and unit variance before concatenating across scans.

Heart rate (HR) and respiration variation (RV) were calculated according to previously described methods (*59*). Before calculating HR and RV, the photoplethysmograph and respiration data accompanying each scan were manually inspected. We selected 104 scans from 26 participants for HR and 51 scans from 31 participants for RV. To calculate the HR, the photoplethysmograph accompanying each scan were first filtered using bandpass filters (1 Hz < f < 150 Hz) to facilitate peak detection. Then, the HR was estimated as the inverse of the average peak-to-peak interval in a 6 s sliding window. RV was calculated as the temporal standard deviation of the raw respiration signal, accompanying each scan, in a 6-second sliding window.

### Autoregressive models (AR) and surrogate data generation

For each sample of real data (single-scan or concatenated scan), we fitted AR models with various model-orders (lags) to rs-fMRI data using the procedure described in a previous study (*17*) and the code distributed by the authors. The lag in AR ranged from 1 to 16. The obtained AR parameters ([A*_1_*… A*_l_*]) were used to generate the surrogate data. For both Gaussian and non-Gaussian AR surrogates, simulated data were generated using a randomly selected time point from the real fMRI data as the seed, and by iteratively applying the autoregressive equation with Gaussian or non-Gaussian noise, respectively. The noise for the Gaussian AR surrogate was drawn from zero-mean multivariate Gaussian noise with a covariance matrix matching to that of the AR residual. The noise for the non-Gaussian AR surrogate is drawn from the temporally shuffled AR residual. The noise for the block-wise non-Gaussian AR surrogate (Fig. 2A) was obtained by first dividing the AR residual into blocks and then temporally shuffling the blocks.

In addition to AR-based surrogates, surrogate data based on phase randomization (PR) were also used. PR retains the complete autoregressive structures and the covariance structures of the real data (*17*). PR surrogates were generated by applying the discrete Fourier transform (DFT) to the real rs-fMRI data. Random phases were then added to the Fourier-transformed data, and the inverse DFT was applied. The added phases were independently generated for each frequency but were the same across brain regions (*17*).

### Innovation driven coactivation pattern analysis (iCAP)

We focused on innovation signal because previous studies have shown that their distribution differs between real rs-fMRI data and surrogate data generated by PR (*13, 21*). The analysis was conducted following the procedures described by Karahanoglu et al., with MATLAB codes provided by the researchers (*13, 60*). Activity-inducing signal and innovation signal were obtained by deconvolving real or surrogate rs-fMRI data using the total activation framework (*60*). Temporal regularization was imposed so that the activity-inducing signal showed block-type activation patterns (without prior knowledge of the duration and timing of the blocks). Because the analysis was not voxel-based, we did not impose a spatial smoothness constraint. For each time course of the innovation signal, the first ten volumes were discarded because the first few volumes tended to show artifacts related to the start of the time course. Dunn’s test, corrected by the Bonferroni method, was applied to the kurtosis of the distribution of innovation signals to assess the statistical significance of the difference between real and surrogate data.

### Topological data analysis (TDA)

We conducted a Mapper-based TDA following the procedures described by Saggar et al. with the MATLAB codes in their study (*33*). Briefly, in the first step, high-dimensional input data were embedded in a two-dimensional space using a filter function. To capture the intrinsic geometry of the data, we used a nonlinear filter function based on neighborhood embedding. Specifically, Euclidian distances were calculated for all pairs of volumes. A k-nearest neighbor graph was then constructed using all the volumes and calculated distances. Using the k-nearest neighbor graph, the geodesic distances were calculated for all volumes in the input space. The geodesic distance was then embedded into a two-dimensional Euclidean space using multi-dimensional scaling. In the second step, overlapping two-dimensional binning was performed for data compression and noise reduction. Based on a previous study by Saggar *et al.* (*33*), we chose a resolution parameter of 14. In the third step, partial clustering was performed within each bin. In the final step, a shape-graph was generated by connecting the nodes from different bins when any volumes were shared by the bins. For statistical comparison, as in a previous study (*33*), we calculated the proportion of high-degree nodes (> 20 degrees). The statistical significance of the differences between the real and surrogate data was assessed using one-way analysis of variance (ANOVA). We also applied Dunn’s test with Bonferroni correction to the number of nodes in each graph to assess the statistical significance of the difference between the real and surrogate data.

### Analyses of AR-10 residuals

To obtain normalized residuals, PCA was applied to AR-10 residuals to spatially decorrelate the signal. After standardizing each PC zero mean and unit variance, time courses of all PCs were concatenated to yield normalized residuals. An analysis of the non-stationarity of the scan-concatenated data was performed by applying PCA and ICA to the residuals of AR-10 fitted to the scan-concatenated data. The ICA was conducted using the MATLAB function rica with 15 ICs. All the time courses of the 15 ICs from all 103 participants were clustered into 15 clusters using k-means clustering. The time course of each cluster was obtained by averaging the ICs in each cluster.

Residual power was obtained by first squaring AR-10 residuals and then taking the 1^st^ PC. Residual power was then compared with non-neuronal variables (HR, RV and FD) using Pearson correlation.

### Data and code availability

The data used in this study are available from the website of (*51*) or from HCP. All essential codes for reproducing the results will be made publicly available upon publication.

## Author Contributions

**TM:** Conceptualization, Formal analysis, Investigation, Visualization, Writing - Review & Editing. **RL:** Investigation, Writing - Review & Editing. **KM:** Formal analysis, Investigation, Writing - Review & Editing. **KJ:** Investigation, Data Curation, Writing - Review & Editing.

## Acknowledgments

Data provided in part by the Human Connectome Project, WU-Minn Consortium (Principal Investigators: David Van Essen and Kamil Ugurbil; 1U54MH091657) funded by the 16 NIH Institutes and Centers that support the NIH Blueprint for Neuroscience Research; and by the McDonnell Center for Systems Neuroscience at Washington University.

## Data and Code Availability

The data used in this study are available from HCP. All codes used for the analysis will be made available upon acceptance.

## Supplementary Figures

**Figure S1.**
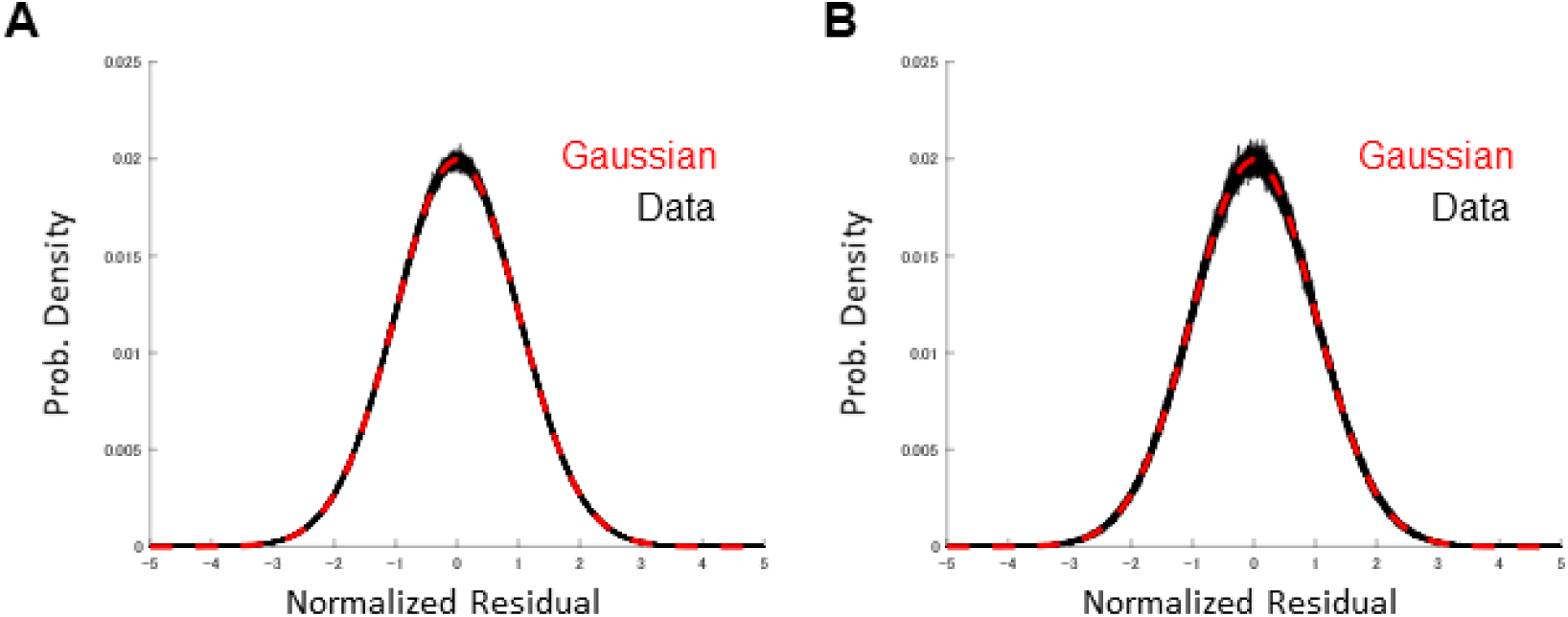
Same as Figure 2F and 4F but for PR surrogates. **(A)** Same plot as Figure 2F, but for PR surrogates of scan-concatenated data. **(B)** Same plot as Figure 4F, but for PR surrogates of single-scan data.

**Figure S2.**
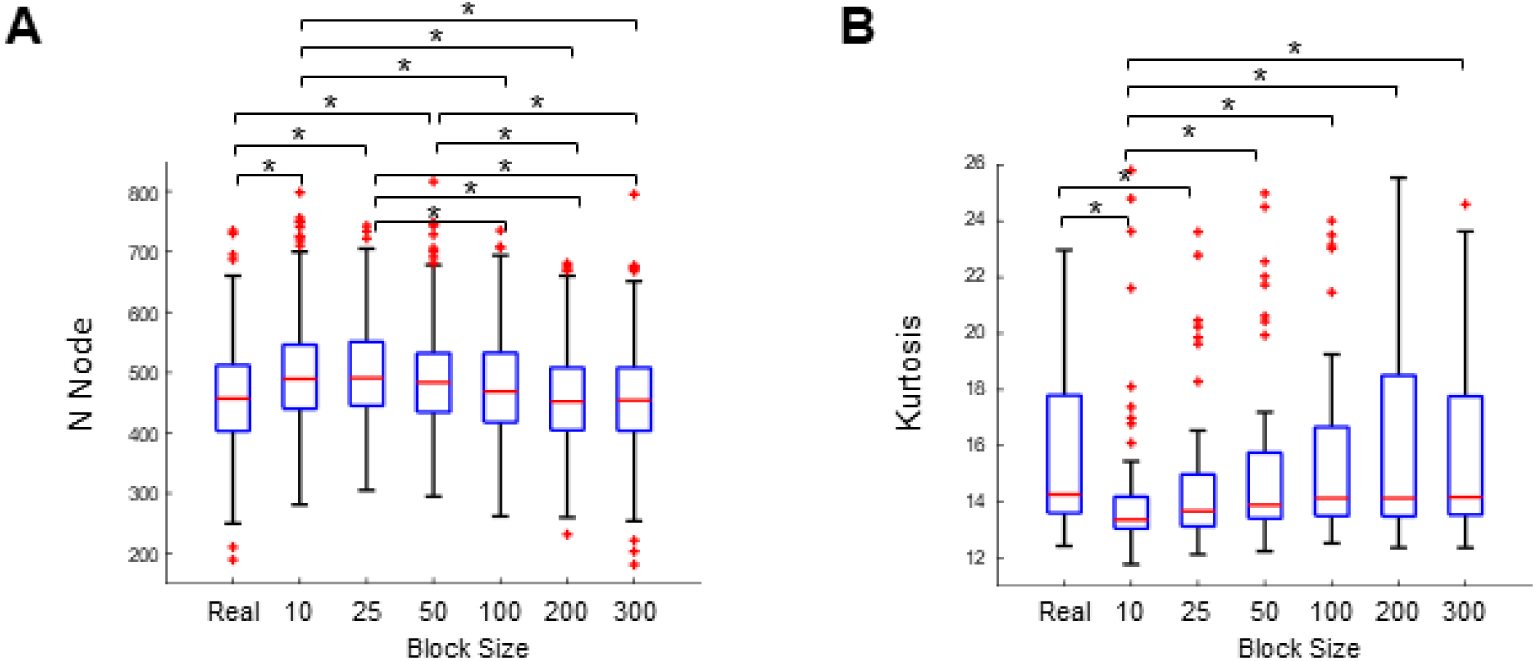
Same as Figure 5C and E but significant differences are indicated. **(A)** Same plot as Figure 5C, but significant differences are indicated. **(B)** Same plot as Figure 5E, but significant differences are indicated. (i.e., excluding interpolated frames). *, p < 0.05 (Dunn’s test, Bonferroni corrected).

**Figure S3.**
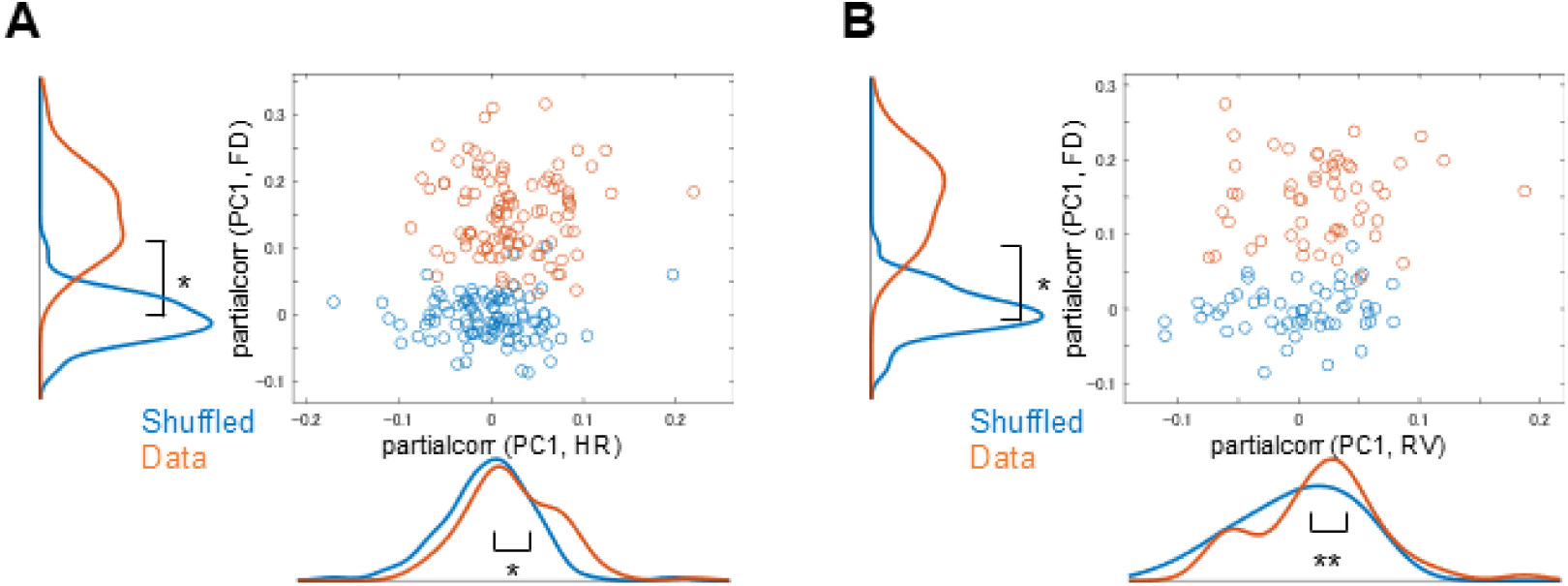
Same as Figure 6B and C but calculated using only censored frames. **(A)** Scatter plot showing correlation between PC1 and HR (FD) partialized for FD (HR) for all scans selected for good HR measurement (104 scans). Same convention as Figure 6B, calculated using only censored frames (i.e., excluding interpolated frames). **(B)** Scatter plot showing the correlation between PC1 and RV (FD) partialized for FD (HR) for all scans selected for good RV measurement (51 scans). Same convention as Figure 6C, calculated using only censored frames (i.e., excluding interpolated frames). *, p < 0.001, **, p < 0.05 (two-sample t test).

## References

1. R. M. Hutchison et al., Dynamic functional connectivity: promise, issues, and interpretations. Neuroimage 80, 360–378 (2013).

2. M. G. Preti, T. A. Bolton, D. Van De Ville, The dynamic functional connectome: State-of-the-art and perspectives. Neuroimage, (2016).

3. V. D. Calhoun, R. Miller, G. Pearlson, T. Adalı, The chronnectome: time-varying connectivity networks as the next frontier in fMRI data discovery. Neuron 84, 262–274 (2014).

4. J. Faskowitz, F. Z. Esfahlani, Y. Jo, O. Sporns, R. F. Betzel, Edge-centric functional network representations of human cerebral cortex reveal overlapping system-level architecture. Nat Neurosci 23, 1644–1654 (2020).

5. T. Bolt et al., A parsimonious description of global functional brain organization in three spatiotemporal patterns. Nat Neurosci 25, 1093–1103 (2022).

6. J. Cabral, F. F. Fernandes, N. Shemesh, Intrinsic macroscale oscillatory modes driving long range functional connectivity in female rat brains detected by ultrafast fMRI. Nat Commun 14, 375 (2023).

7. J. C. Pang et al., Geometric constraints on human brain function. Nature 618, 566–574 (2023).

8. C. Chang, G. H. Glover, Time-frequency dynamics of resting-state brain connectivity measured with fMRI. Neuroimage 50, 81–98 (2010).

9. T. Watanabe, G. Rees, Brain network dynamics in high-functioning individuals with autism. Nat Commun 8, 16048 (2017).

10. R. V. Raut et al., Global waves synchronize the brain’s functional systems with fluctuating arousal. Sci Adv 7, (2021).

11. E. C. Hansen, D. Battaglia, A. Spiegler, G. Deco, V. K. Jirsa, Functional connectivity dynamics: modeling the switching behavior of the resting state. Neuroimage 105, 525–535 (2015).

12. D. Vidaurre, S. M. Smith, M. W. Woolrich, Brain network dynamics are hierarchically organized in time. Proc Natl Acad Sci U S A 114, 12827–12832 (2017).

13. F. I. Karahanoğlu, D. Van De Ville, Transient brain activity disentangles fMRI resting-state dynamics in terms of spatially and temporally overlapping networks. Nat Commun 6, 7751 (2015).

14. X. Liu, J. H. Duyn, Time-varying functional network information extracted from brief instances of spontaneous brain activity. Proc Natl Acad Sci U S A 110, 4392–4397 (2013).

15. D. J. Lurie et al., Questions and controversies in the study of time-varying functional connectivity in resting fMRI. Netw Neurosci 4, 30–69 (2020).

16. T. O. Laumann et al., On the Stability of BOLD fMRI Correlations. Cereb Cortex, (2016).

17. R. Liégeois, T. O. Laumann, A. Z. Snyder, J. Zhou, B. T. T. Yeo, Interpreting temporal fluctuations in resting-state functional connectivity MRI. Neuroimage 163, 437–455 (2017).

18. T. Matsui, K. I. Yamashita, Static and Dynamic Functional Connectivity Alterations in Alzheimer’s Disease and Neuropsychiatric Diseases. Brain Connect 13, 307–314 (2023).

19. D. Vidaurre, Dynamic functional connectivity: Why the controversy?. Imaging Neuroscience 2, 1–4 (2024).

20. T. O. Laumann, A. Z. Snyder, C. Gratton, Challenges in the measurement and interpretation of dynamic functional connectivity. Imaging Neuroscience 2, 19 (2024).

21. D. Van De Ville, R. Liégeois, Dynamic functional connectivity to tile the spatiotemporal mosaic of brain states. Imaging Neuroscience 2, 1–5 (2024).

22. E. Tagliazucchi, Rest assured: Dynamic functional connectivity and the baseline state of the human brain. Imaging Neuroscience 2, 1–7 (2024).

23. X. Liu, C. Chang, J. H. Duyn, Decomposition of spontaneous brain activity into distinct fMRI co-activation patterns. Front Syst Neurosci 7, 101 (2013).

24. T. Matsui, T. Q. Pham, K. Jimura, J. Chikazoe, On co-activation pattern analysis and non-stationarity of resting brain activity. Neuroimage 249, 118904 (2022).

25. Y. Hosaka et al., Surrogate data analyses of the energy landscape analysis of resting-state brain activity. Front Neural Circuits 19, 1500227 (2025).

26. Z. Ladwig et al., BOLD cofluctuation ‘events’ are predicted from static functional connectivity. Neuroimage 260, 119476 (2022).

27. T. Matsui, T. Murakami, K. Ohki, Neuronal Origin of the Temporal Dynamics of Spontaneous BOLD Activity Correlation. Cereb Cortex, (2018).

28. T. O. Laumann, A. Z. Snyder, C. Gratton, Challenges in the measurement and interpretation of dynamic functional connectivity. Imaging Neuroscience 2, 19 (2024).

29. F. Freyer, K. Aquino, P. A. Robinson, P. Ritter, M. Breakspear, Bistability and non-Gaussian fluctuations in spontaneous cortical activity. J Neurosci 29, 8512–8524 (2009).

30. J. A. Roberts, T. W. Boonstra, M. Breakspear, The heavy tail of the human brain. Curr Opin Neurobiol 31, 164–172 (2015).

31. F. de Pasquale et al., Temporal dynamics of spontaneous MEG activity in brain networks. Proc Natl Acad Sci U S A 107, 6040–6045 (2010).

32. R. F. Betzel et al., Synchronization dynamics and evidence for a repertoire of network states in resting EEG. Front Comput Neurosci 6, 74 (2012).

33. M. Saggar, J. M. Shine, R. Liégeois, N. U. F. Dosenbach, D. Fair, Precision dynamical mapping using topological data analysis reveals a hub-like transition state at rest. Nat Commun 13, 4791 (2022).

34. A. Schaefer et al., Local-Global Parcellation of the Human Cerebral Cortex from Intrinsic Functional Connectivity MRI. Cereb Cortex 28, 3095–3114 (2018).

35. E. Nozari et al., Macroscopic resting-state brain dynamics are best described by linear models. Nat Biomed Eng 8, 68–84 (2024).

36. C. Gorrostieta, M. Fiecas, H. Ombao, E. Burke, S. Cramer, Hierarchical vector auto-regressive models and their applications to multi-subject effective connectivity. Front Comput Neurosci 7, 159 (2013).

37. M. Bianciardi et al., Sources of functional magnetic resonance imaging signal fluctuations in the human brain at rest: a 7 T study. Magn Reson Imaging 27, 1019–1029 (2009).

38. T. Bolt et al., Autonomic physiological coupling of the global fMRI signal. Nat Neurosci 28, 1327–1335 (2025).

39. C. Chang et al., Tracking brain arousal fluctuations with fMRI. Proc Natl Acad Sci U S A 113, 4518–4523 (2016).

40. C. Stringer et al., Spontaneous behaviors drive multidimensional, brainwide activity. Science 364, 255 (2019).

41. S. Musall, M. T. Kaufman, A. L. Juavinett, S. Gluf, A. K. Churchland, Single-trial neural dynamics are dominated by richly varied movements. Nat Neurosci 22, 1677–1686 (2019).

42. C. Chang et al., Association between heart rate variability and fluctuations in resting-state functional connectivity. Neuroimage 68, 93–104 (2013).

43. J. D. Power, K. A. Barnes, A. Z. Snyder, B. L. Schlaggar, S. E. Petersen, Spurious but systematic correlations in functional connectivity MRI networks arise from subject motion. Neuroimage 59, 2142–2154 (2012).

44. D. A. Fair et al., Correction of respiratory artifacts in MRI head motion estimates. Neuroimage 208, 116400 (2020).

45. J. Piccinini et al., Data-driven discovery of canonical large-scale brain dynamics. Cereb Cortex Commun 3, tgac045 (2022).

46. A. Kashyap et al., Using an ordinary differential equation model to separate rest and task signals in fMRI. Nat Commun 16, 7128 (2025).

47. C. C. Chen, C. W. Tyler, H. A. Baseler, Statistical properties of BOLD magnetic resonance activity in the human brain. Neuroimage 20, 1096–1109 (2003).

48. T. Okuno et al., Group Surrogate Data Generating Models and similarity quantification of multivariate time-series: A resting-state fMRI study. Neuroimage 279, 120329 (2023).

49. R. Hindriks et al., Can sliding-window correlations reveal dynamic functional connectivity in resting-state fMRI? Neuroimage 127, 242–256 (2016).

50. R. Liégeois et al., Resting brain dynamics at different timescales capture distinct aspects of human behavior. Nat Commun 10, 2317 (2019).

51. T. Ezaki, T. Watanabe, M. Ohzeki, N. Masuda, Energy landscape analysis of neuroimaging data. Philos Trans A Math Phys Eng Sci 375, (2017).

52. A. Xifra-Porxas, M. Kassinopoulos, G. D. Mitsis, Physiological and motion signatures in static and time-varying functional connectivity and their subject identifiability. Elife 10, (2021).

53. R. V. Raut et al., Arousal as a universal embedding for spatiotemporal brain dynamics. Nature 647, 454–461 (2025).

54. R. Li, K. Ohki, T. Matsui, Ketamine-induced 1-Hz oscillation of spontaneous neural activity is not directly visible in the hemodynamics. Biochem Biophys Res Commun 678, 102–108 (2023).

55. J. Cabral et al., Cognitive performance in healthy older adults relates to spontaneous switching between states of functional connectivity during rest. Sci Rep 7, 5135 (2017).

56. R. Chen, M. Singh, T. S. Braver, S. Ching, Dynamical models reveal anatomically reliable attractor landscapes embedded in resting-state brain networks. Imaging Neurosci (Camb) 3, (2025).

57. M. F. Glasser et al., The minimal preprocessing pipelines for the Human Connectome Project. Neuroimage 80, 105–124 (2013).

58. B. T. Yeo et al., The organization of the human cerebral cortex estimated by intrinsic functional connectivity. J Neurophysiol 106, 1125–1165 (2011).

59. C. Chang, G. H. Glover, Relationship between respiration, end-tidal CO2, and BOLD signals in resting-state fMRI. Neuroimage 47, 1381–1393 (2009).

60. F. I. Karahanoğlu, C. Caballero-Gaudes, F. Lazeyras, D. Van de Ville, Total activation: fMRI deconvolution through spatio-temporal regularization. Neuroimage 73, 121–134 (2013).

